# Essential role of the mouse synapse associated protein Syap1 in circuits for spontaneous motor activity and rotarod balance

**DOI:** 10.1101/542829

**Authors:** Cora R. von Collenberg, Dominique Schmitt, Thomas Rülicke, Michael Sendtner, Robert Blum, Erich Buchner

**Affiliations:** Institute of Clinical Neurobiology, University Hospital Würzburg, Versbacher Str. 5, 97078 Würzburg, Germany; Institute of Laboratory Animal Science, University of Veterinary Medicine Vienna, 1210 Vienna, Austria

**Keywords:** *Syap1* knockout, motor behaviour, associative learning, fear conditioning, object recognition

## Abstract

Synapse-associated protein 1 (Syap1) is the mammalian homologue of synapse-associated protein of 47 kDa (Sap47) in *Drosophila*. Genetic deletion of Sap47 leads to deficiencies in short-term plasticity and associative memory processing in flies. In mice, Syap1 is prominently expressed in the nervous system, but its function is still unclear. We have generated *Syap1* knockout mice and tested motor behaviour and memory. These mice are viable and fertile but display distinct deficiencies in motor behaviour. Locomotor activity specifically appears to be reduced in early phases when voluntary movement is initiated. On the rotarod, a more demanding motor test involving control by sensory feedback, Syap1-deficient mice dramatically fail to adapt to accelerated speed or to a change in rotation direction. Syap1 is highly expressed in cerebellar Purkinje cells and cerebellar nuclei. Thus, this distinct motor phenotype could be due to a so far unknown function of Syap1 in cerebellar sensorimotor control. The observed motor defects are highly specific since other tests in the modified SHIRPA test, as well as cognitive tasks like novel object recognition, Pavlovian fear conditioning, and anxiety-like behaviour in open field, dark-light transition, and elevated plus maze, do not appear to be strongly affected in *Syap1* knockout mice.

**Summary statement:** Knockout of the *Syap1* gene in mice causes a distinct motor behaviour phenotype.

## Introduction

Synapse associated protein 1 (Syap1) is a member of synapse associated BSD domain protein family (Doerks et al., 2002). It has been discovered by characterizing antigens using a library of monoclonal antibodies against *Drosophila* head homogenates (Reichmuth et al., 1995; Hofbauer et al., 2009). One of these antibodies binds to fly neuropil and in particular to presynaptic boutons of glutamatergic larval motoneurons. In head homogenate, the antibody detects a protein of 47 kDa, which was termed synapse-associated protein of 47 kDa (Sap47). Cloning and subsequent genetic deletion of the *Sap47* gene revealed that Sap47 is not required for viability. However, knockout larvae showed defects in short-term plasticity and olfactory associative learning and memory (Saumweber et al., 2011). The human gene encoding the synapse-associated protein 1 (*SYAP1*) is located within chromosomal band Xp22.2, a region associated with mental retardation, developmental delay and autism spectrum disorder (Sismani et al., 2011; Prasad et al., 2012).

The function of mammalian Syap1 is largely unclear. Syap1 has been shown to be important for adipocyte differentiation from murine embryonic stem cells by stimulating phosphorylation of Akt1 kinase (Yao et al., 2013). In cultured mouse motoneurons, however, no change in Akt phosphorylation after *Syap1* knockout or knockdown has been observed (Schmitt et al., 2016). Furthermore, size and body weight of *Syap1* knockout male mice (Y/-) are undistinguishable to that of wildtype littermates (Y/+) (Schmitt et al., 2016). Thus, the lack of Syap1 in mice apparently does not impair general metabolism or lipid storage *in vivo*.

The distribution of Syap1 immunoreactivity in the mouse brain has recently been reported (Schmitt et al., 2016). Syap1 is widely found throughout synaptic neuropil with high concentrations in regions rich in glutamatergic synapses. In addition, it has been detected in perinuclear structures in close proximity to the Golgi apparatus of subgroups of neurons, suggesting that it may be involved in a more general process of vesicular trafficking in addition to its putative role in synaptic transmission.

Here we describe the first behavioural analysis of *Syap1* knockout mice. Our data show that male mice hemizygous for a *Syap1* null allele (*Syap*^*Y/-*^) exhibit a distinct motor phenotype in the open field test and on the rotarod during change of speed or direction. The main phenotype of the mutants might point to a specific so far unknown function of Syap1 in cerebellar circuits for motor control. These findings could be of clinical relevance for patients with mutations in the *SYAP1* gene on chromosome Xp22.2.

## Results

Observation of voluntary behaviour and movements of *Syap1* knockout animals in their cages revealed that knockout males, compared to wildtype littermates, show less exploratory activity and require longer to restore normal motor activity post handling. We therefore performed standardized behavioural phenotyping of these animals. We conducted a modified SHIRPA test (summarized in table 1) covering 20 measures of sensorimotor functions and reflexes (Rogers et al., 1997, Hatcher et al., 2001). Most aspects of the behaviour of *Syap1* knockout animals were inconspicuous in comparison to their wildtype littermates, indicating that lack of Syap1 does not cause generalized neurological defects. However, *Syap1*^*Y/-*^ mice showed a specific deficit in locomotor activity, assessed by the number of squares crossed within a given time (table 1).

**Table 1:** Modified SHIRPA test results

| **SHIRPA** |  |  | *Syap1*<sup>Y/+</sup> | *Syap1*<sup>Y/-</sup> |
| --- | --- | --- | --- | --- |
| ***viewing jar*** | body position | active | 12 / 12 | 12 / 12 |
|  |  | inactive | 0 / 12 | 0 / 12 |
|  | tremor | present | 0 / 12 | 0 / 12 |
|  | palpebral closure | eyes open | 12 / 12 | 12 / 12 |
|  | coat appearance | well groomed | 12 / 12 | 12 / 12 |
|  | whisker | intact | 12 / 12 | 12 / 12 |
|  | lacrimation | present | 0 / 12 | 0 / 12 |
|  | defecation | present | 0 / 12 | 1 / 12 |
|  | urination | present | 0 / 12 | 0 / 12 |
| ***arena*** | transfer arousal | immediate movement | 12 / 12 | 12 / 12 |
|  | **locomotor activity\*** | squares crossed in 30 s | 10,42 | 6,92 |
|  | gait | fluid movement | 12 / 12 | 12 / 12 |
|  | tail elevation | elevated tail | 0 / 12 | 0 / 12 |
|  | touch escape | response to touch | 11 / 12 | 12 / 12 |
|  |  | flees prior to touch | 1 / 12 | 0 / 12 |
| ***above arena*** | skin colour | pink | 12 / 12 | 12 / 12 |
|  | trunk curl | present | 0 / 12 | 0 / 12 |
|  | limp grasping | present | 0 / 12 | 0 / 12 |
|  | pinna reflex | present | 12 / 12 | 12 / 12 |
|  | corneal reflex | present | 12 / 12 | 12 / 12 |
|  | evidence of biting | present | 0 / 12 | 0 / 12 |
|  | vocalization | present | 0 / 12 | 0 / 12 |

\* unpaired t-test: **p = 0.0134**

The SHIRPA data indicated a defect in motor behaviour and therefore we first performed the open field test (OF) to investigate the overall locomotor activity of *Syap1*-deficient mice (Hall, 1934, Carola et al., 2002). Notably, *Syap1* knockout mice showed a reduction in the total travel distances within the first 10 min of the OF test (Fig. 1A). This reduction was more pronounced in the periphery (Fig. 1C) than in the centre (Fig. 1B). *Syap1* knockout mice were more or less stationary during the first five minutes of the OF, before they regained their normal locomotor activity. Orientation-like movements on the same spot characterized these atypical early hypoactive phases. The atypical motor behaviour of *Syap1* knockout mice in the open field test is shown in the supplementary movies (*Syap1*^*Y/-*^: supplementary movie S1; wildtype: supplementary movie S2). This behaviour was clearly distinguishable from freezing behaviour, a typical defensive reaction of mice (Tovote et al., 2015). In the movement analysis, wildtype mice exhibit an overall mean speed of 7.7 cm/s, while *Syap1* knockout mice did not reach this mean speed (indicated as a red horizontal line in Fig. 1D,F,H,J). In later phases, after 15 – 30 minutes in the test arena, this phenotype was lost (Fig. 1A). We also plotted the locomotor speed over time to better illustrate this main effect (Fig. 1 D-K). In later phases of the test, from minute 15 – 30, *Syap1* knockout mice show the capability to walk at similar speed as wildtype littermates (Fig. 1D-K). Tracking for locomotor activity reveals that the *Syap1* knockout phenotype is mainly caused by the typical longer stationary phases (red labels in Fig. 1E,G,I,K). Significant differences in movement speed between the genotypes arose within the first 10 min of the OF (supplementary Fig. S1A).

**Figure 1:**
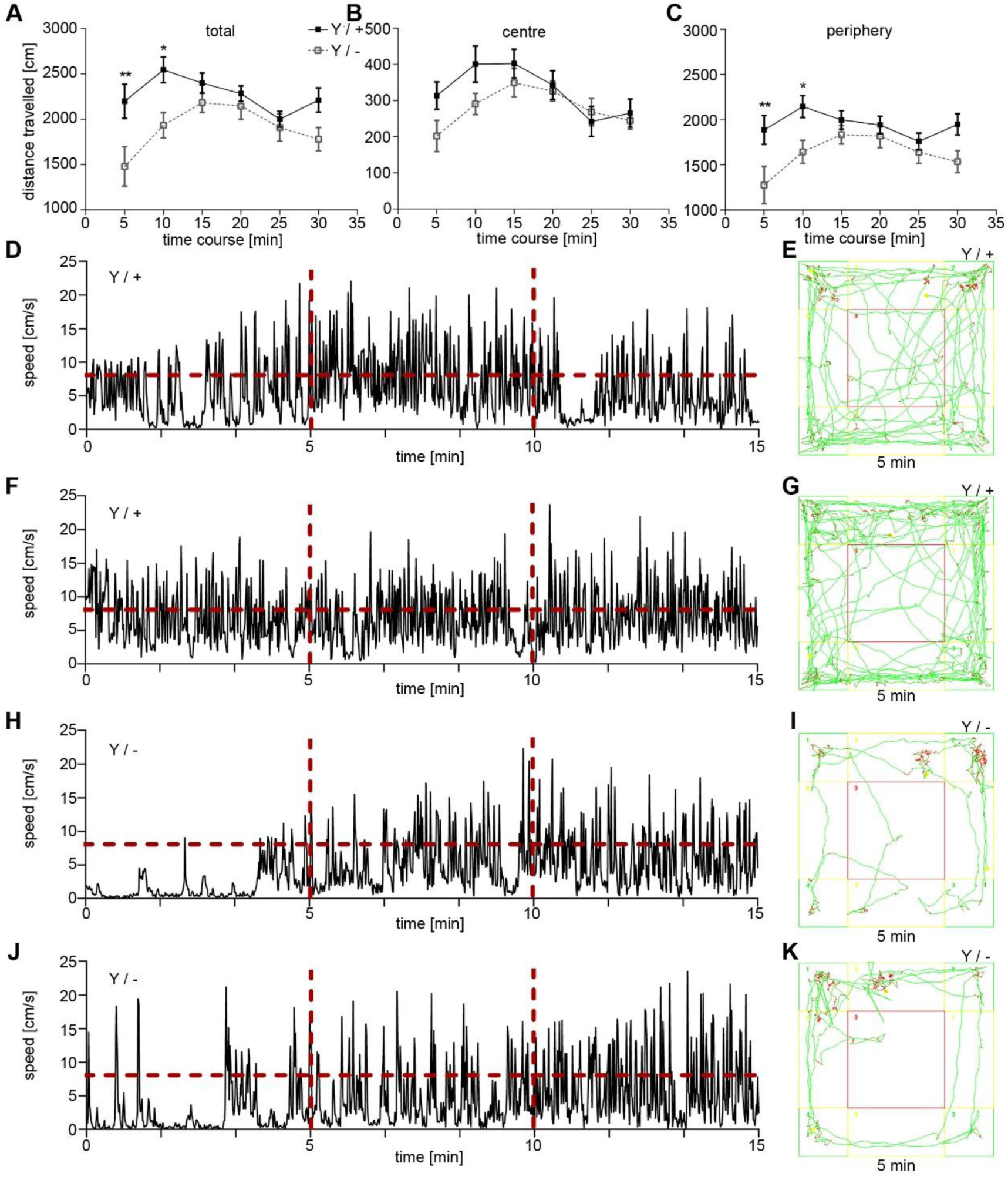
Reduced locomotor activity of male *Syap1* knockout mice in early phases of the open field test. (A) Total distance travelled per 5 min in the open field test. *Syap1* knockout animals travelled less within the first 10 min of the open field test (A, Sidak’s multiple comparison: * p = 0.0164, ** p = 0.0027, all other time points p > 0.05). (B, C) Total distance travelled per 5 min periods in the centre and periphery of the open field test (Sidak’s multiple comparison: * p = 0.04, ** p = 0.0063). (D, F, H, J) Representative graphs showing the speed of movement for two wildtype and two *Syap1* knockout mice. The horizontal red line indicates the mean overall speed of movement of wildtype mice (7.7 cm/s). (E, G, I, K) Tracks showing the locomotor behaviour in the OF arena within the first five minutes of the test for wildtype (in E, G) and *Syap1* knockout mice (in I, K), respectively. Red colour indicates movement at a speed of less than 4.81 cm/s. Further statistical values are given in supplementary table S1.

The OF is also suited to examine anxiety-like behaviour in rodents (Hall, 1934, Carola et al., 2002). In contrast to the considerable changes in early locomotor activity, the time spent in the centre of the open field was not significantly different between the genotypes (supplementary Fig. S1B,C). The initial difference in the number of entries into centre and periphery (supplementary Fig. S1D,E) was most likely a consequence of the reduced initial locomotor activity.

A stringent test on motor coordination under proprioceptive feedback control is obtained when instead of walking on a flat surface the animal has to maintain its balance on a rotating rod (rotarod) (Jones and Roberts, 1968, Shiotsuki et al., 2010). Under condition of steady rotation, *Syap1* knockout mice, like wildtype mice, managed to stay and walk on the rod for the entire observation period of 5 min (Fig. 2A). However, in more challenging situations, the *Syap1* mutants show clear deficiencies, like falling off the rod with very short latency when rod rotation was accelerated or reversed (rocking) (Fig. 2B,C). In both cases, the differences between the genotypes are highly significant (accelerated: p < 0.0001; rocking: p < 0.0001, two-way ANOVA). It is very unlikely that this effect was due to general muscle weakness as grip strength was only slightly impaired in the hind limbs of the mutants (Fig. 2D, Sidak’s multiple comparisons: p = 0.0041). We also tested whether *Syap1*^*Y/-*^ mice take advantage of voluntary running in a wheel. Notably, voluntary running distance over night was not reduced in *Syap1* knockout mice compared to wildtype littermates (*Syap* ^*Y/-*^: 3.5 km; wildtype: 3.6 km; mean of n = 3 per genotype).

**Figure 2:**
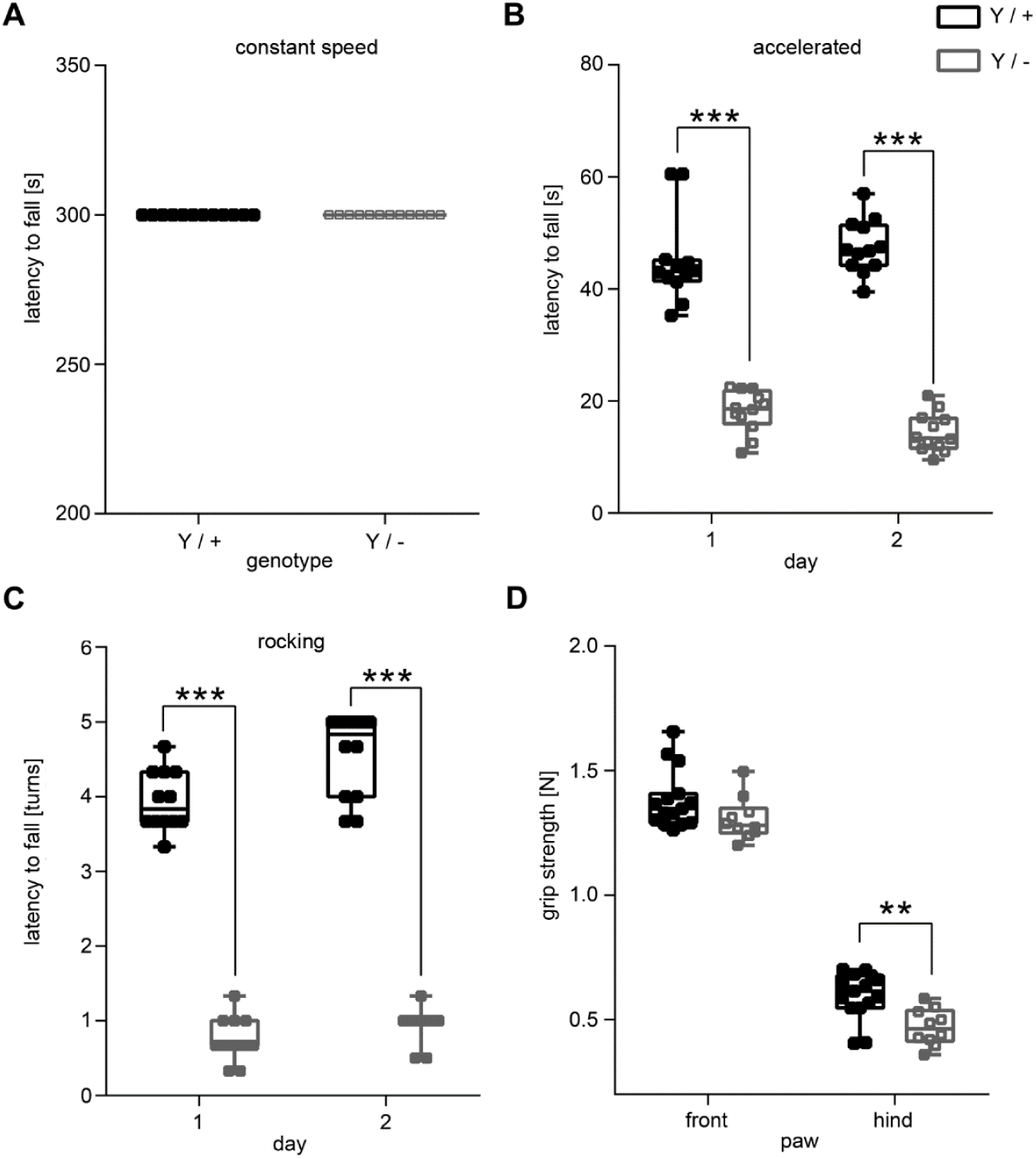
Male *Syap1* knockout mice fail in the accelerated and rocking rotarod test. (A) All animals managed to stay on a rod rotating at constant speed (5 rpm) for 5 min. (B, C) When rotation was accelerated (in B) or reversed (in C, rocking), latency to fall from the test apparatus was dramatically reduced in Syap1 mutants (B, C, Sidak’s multiple comparison: *** p < 0.001). (D) Grip strength of the hind limb was slightly, but significantly reduced in *Syap1* knockout mice (Sidak’s multiple comparison: ** p <= 0.0041). Further statistical values: supplementary table S1.

In *Drosophila*, knockout of the *Syap1* homologue *Sap47* causes impaired synaptic plasticity and reduced associative olfactory learning (Saumweber et al., 2011). We therefore investigated memory-related behaviour in *Syap1* knockout mice. We first performed the novel object recognition (NOR) test (Ennaceur and Delacour, 1988, Antunes and Biala, 2012). In the NOR test, the performance of the animals primarily relies on innate exploratory behaviour and does not involve reinforcement procedures such as food reward or electric foot shocks. In this task, a novel object needs to be noticed and processed as a memory trace. Furthermore, the pre-existing memory trace of the familiar object needs to be recalled after a certain delay (Ennaceur, 2010). Memory consolidation in the NOR test seems to be hippocampus-dependent and involves synaptic plasticity processes in the perirhinal cortex (Rampon et al., 2000, Vignoli et al., 2016, Antunes and Biala, 2012). The test was performed in the open field arena where mice were confronted with two identical objects on the first day (Fig. 3A). On the next day, one of the objects was substituted with a new one to see whether the mice remember the familiar object and spend more time exploring the novel object (Fig. 3A). Fig. 3B shows that *Syap1* knockout as well as the wildtype littermates spent almost equal amounts of time with two identical objects on day 1 (p = 0.1907). On the next day, wildtype and *Syap1*^*Y/-*^ mice both preferred the novel object (p < 0.001), indicating that basic visual pattern recognition and pattern memory is not affected by Syap1 deficiency in mice (Fig. 3C). This shows that working memory and task-specific episodic memory elements are inconspicuous in *Syap1*^*Y/-*^ mice.

**Figure 3:**
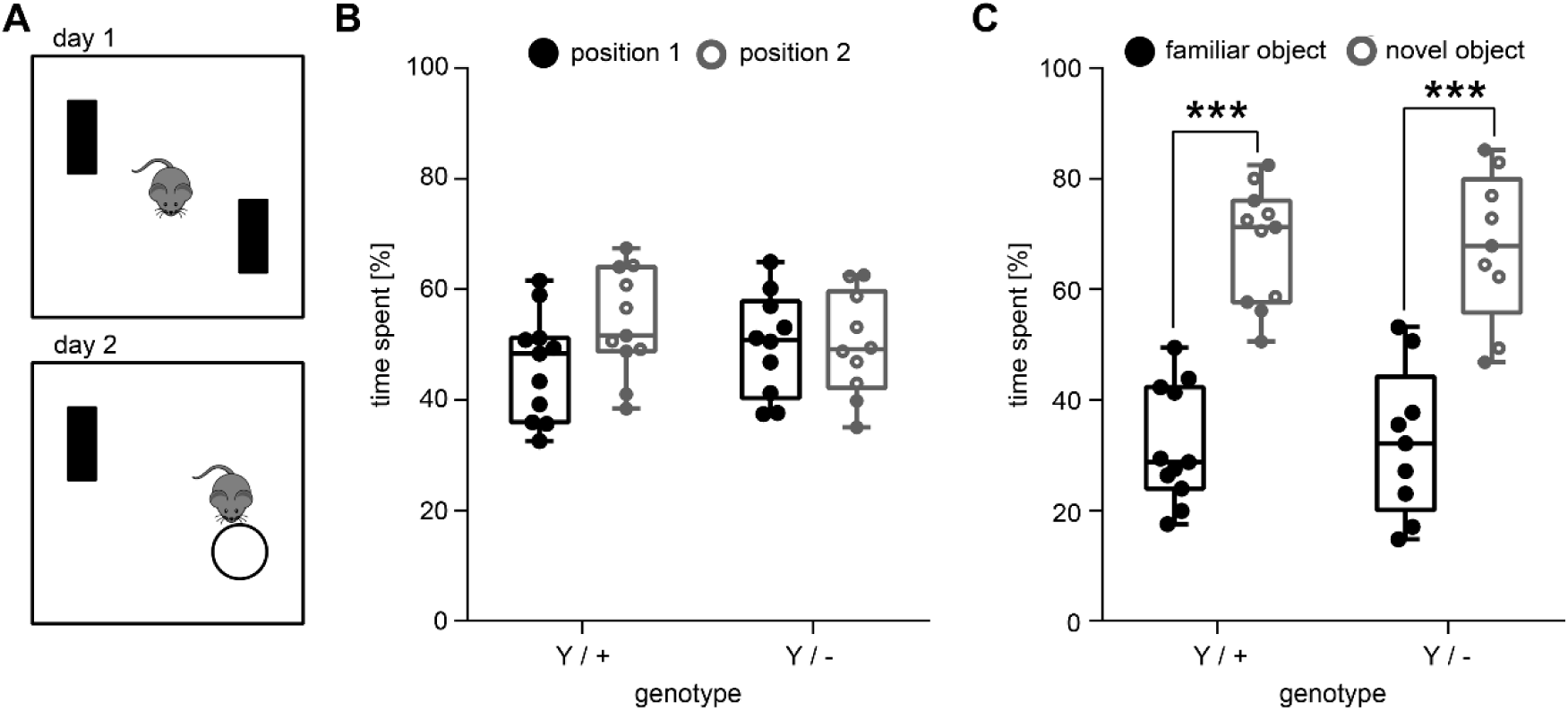
Object recognition memory is not altered in *Syap1* knockout males. (A) New object recognition paradigm. On the first day, the mice were confronted with two identical objects. 24 hours later, the mice were placed back into the same arena, where one of the objects had been substituted with a novel one. (B) *Syap1* knockout and wildtype mice spent the same amount of time at either of two identical objects. Both genotypes spent significantly more time near the novel object on the next day (C, Sidak’s multiple comparison: *** p < 0.0001). The summary of the statistical values is given in supplementary table S1.

We also tested *Syap1* knockout mice in a Pavlovian fear conditioning paradigm (Tovote et al., 2015, Johansen et al., 2011, LeDoux, 2014) that includes both cued and contextual fear conditioning (Fig. 4A). Pavlovian fear conditioning in mice is a typical paradigm to test for associative learning and memory processing (Tovote et al., 2015, Johansen et al., 2011, LeDoux, 2014). Plasticity defects in the amygdala are known to interfere with the conditioning of defensive behaviour to an auditory cue (tone) and to the training context, whereas conditioning to the training context (Context A), but not to the cue, is hippocampus dependent (Maren and Hobin, 2007, Maren et al., 2013, Phillips and LeDoux, 1992). *Syap1* knockout mice displayed freezing behaviour similar to their wildtype littermates in both tests (Fig. 4). The freezing rate increased in both genotypes after initial shock administration indicating inconspicuous fear memory acquisition (Fig. 4B). A significant difference between genotypes (p = 0.0194) was further revealed by the two-way ANOVA. This genotype effect may however be caused by a difference in immobility prior to footshock application (unpaired t-test, p = 0.049). On the next day, when mice were placed in a new context (Context B) but were confronted with the same tone that preceded shock delivery the previous day (recall of tone), both, knockout and wildtype mice, spent significantly more time freezing during tone presentation (Fig. 4C). No significant genotype effect was detected in this case. On the third day, mice were again exposed to the training Context A. Compared to exposure to the neutral Context B on the previous day, the recall of the training context resulted in much higher amount of time spent freezing during Context A exposure (Fig. 4D). Here again a genotype effect was detected which is reflected by longer duration of freezing of the *Syap1* knockout mice in Context A (Fig. 4D, Sidak’s multiple comparison, p = 0.0021). These data suggest that *Syap*^*Y/-*^ mice are able to process associative memory in the fear circuit, as indicated by the cue- and context-dependent elements of the conditioning paradigm. The data also show the ability for context discrimination in *Syap*^*Y/-*^ mice, a function attributed to the hippocampus (Frankland et al., 1998, Maren et al., 2013).

**Figure 4:**
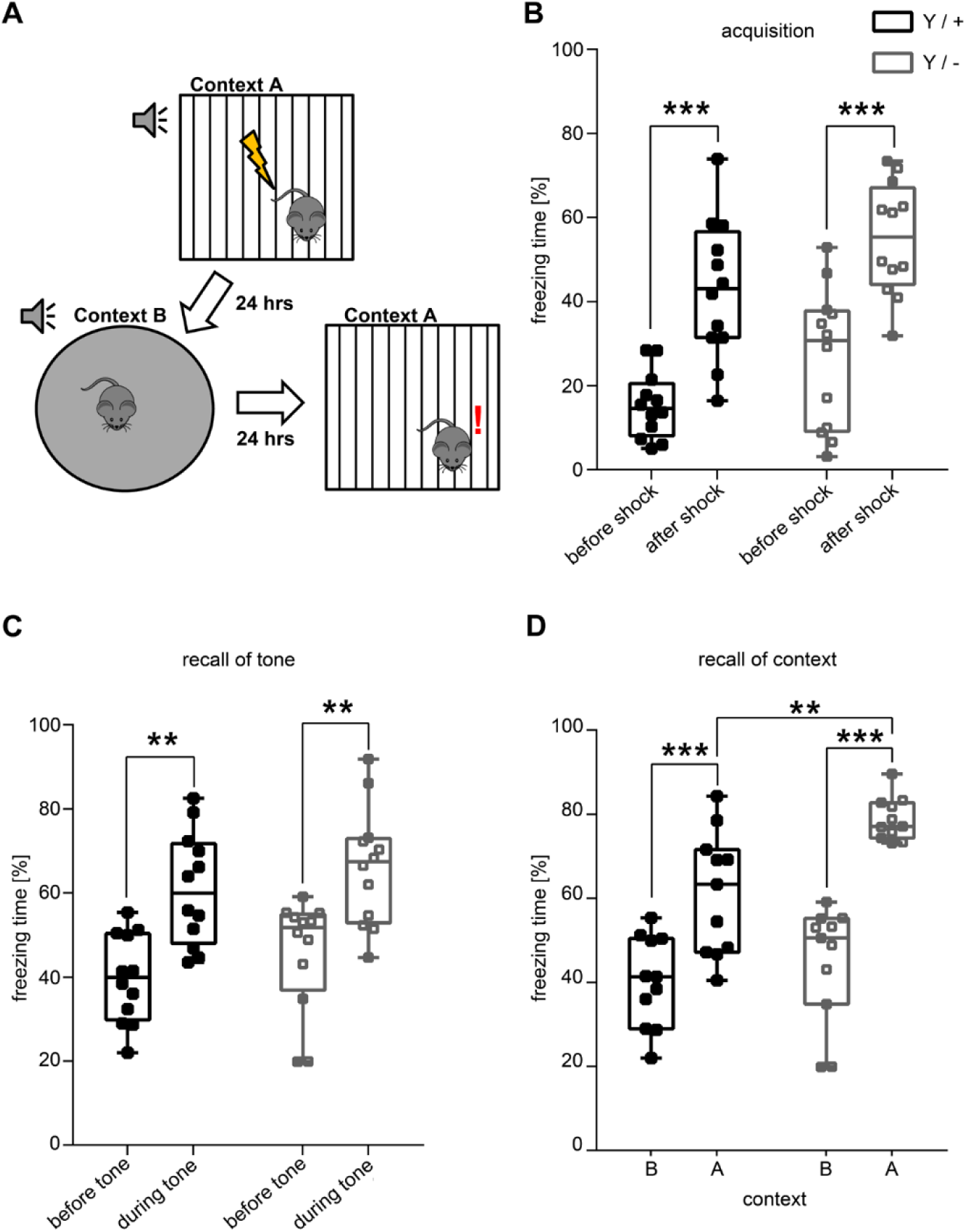
Critical Pavlovian fear conditioning parameters are inconspicuous in *Syap1* knockout mice. (A) Fear conditioning paradigm. Mice were conditioned in Context A, where an acoustic cue was followed by administration of an electric foot shock (unconditioned stimulus). One day later, mice were confronted with the same cue in a novel context (Context B). On the third day, mice were placed in Context A, but without cue presentation. (B) Fear acquisition. Wildtype and *Syap1* knockout mice displayed similar freezing behaviour after shock administration (Sidak’s multiple comparison: *** p < 0.001). (C) Tone recall in Context B. In Context B, both genotypes showed more freezing behaviour during tone presentation (Sidak’s multiple comparison: *** p < 0.001). (D) Context recall. Both genotypes freeze significantly more in Context A than in Context B (D, Sidak’s multiple comparison: ** p = 0.0021, *** p < 0.001). The summary of the statistical values is given in supplementary table S1.

Next, we performed two more tests to investigate explorative activity and anxiety-like behaviour; the dark-light (DL) transition test (Crawley and Goodwin, 1980) and the elevated plus-maze (EPM) test (Pellow et al., 1985, Carola et al., 2002). In a dark-light arena, no differences in the time spent in the illuminated light chamber versus the dark chamber were observed (Fig. 5A). However, as already seen in the SHIRPA and OF tests, reduced locomotor activity was indicated by fewer distances travelled by the *Syap1* mutants (Fig. 5B). In the EPM, independent of the genotype, mice spent more time in the closed arms than in the open arms of the test apparatus (Fig. 5C). Here, the distance travelled was inconspicuous between the genotypes (Fig. 5D). Both groups travelled more in the closed arms of the maze. The DL transition test and the EPM are in accordance with the results obtained in the open field arena (Fig. 1, Fig. S1) and confirm that the reduced motor performance of *Syap*^*Y/-*^ mice is unlikely to be caused by increased anxiety-like behaviour.

**Figure 5:**
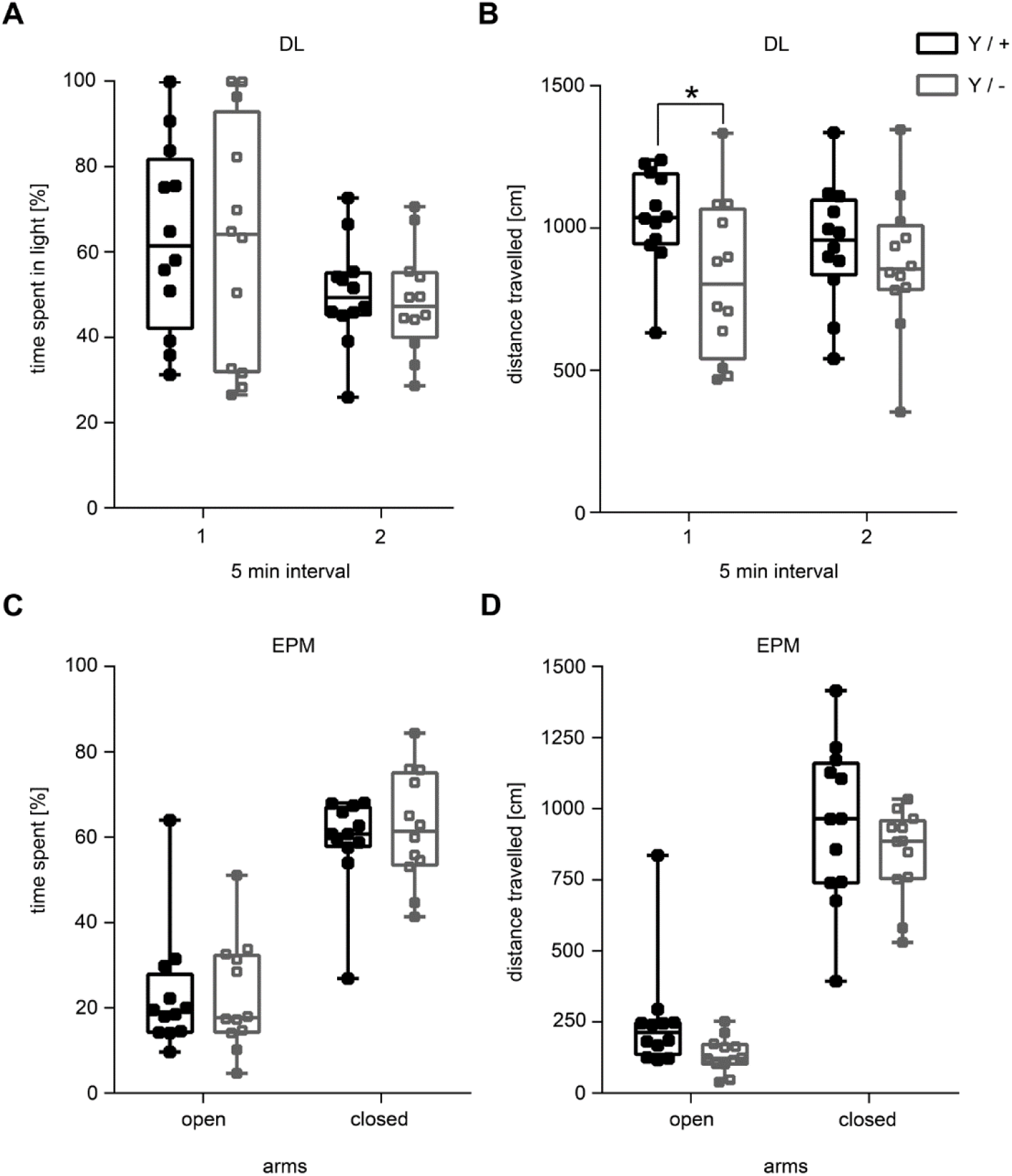
No anxiety-like behaviour in male *Syap1* knockout mice in the dark-light transition and elevated plus maze tests. (A) *Syap1* knockout mice and wildtype littermates spent an equivalent amount of time in the light and dark areas of the arena, indicating no anxiety-like behaviour in this test. (B) Within the first five minutes of the test, the distance travelled in the dark-light arena was slightly reduced in the mutant (Sidak’s multiple comparison: * p = 0.0471). (C) *Syap1* knockout mice and wildtype littermates spent the same amount of time on the open arm of the elevated plus maze. (D) In the elevated plus maze test, total distance travelled did not differ between the two genotypes. The summary of the statistical values is given in supplementary table S1.

## Discussion

Syap1-deficient mice show a distinct locomotor phenotype, whereas in cognition-related behaviour tests they appear inconspicuous. The motor phenotype is most obvious in early phases of the open field and in the accelerated rotarod tests. Interestingly, mice spent more time being stationary in early phases of the open field test, when voluntary initiation of locomotion is needed. Notably, in the late phase of the open field test, Syap1-defienct mice show normal exploratory behaviour, indicating that the mice are able to adapt to the task. *Syap*^*Y/-*^ mice easily manage the rotating rod at a constant speed and take advantage of voluntary wheel running, but they fail in the accelerated or reversed rotarod. The data argue against a fundamental defect in reflexive, rhythmic, and voluntary motor performance or muscle contraction coordination, but point to distinct changes in subcortical motor control centres (Bostan and Strick, 2018, Grillner et al., 2005, De Zeeuw and Ten Brinke, 2015, Calabresi et al., 2014, Kiehn, 2016). Furthermore, the phenotype seems to be different from deficiencies observed in genetic or induced mouse models of Parkinson’s disease, where subtle alterations in the nigrostriatal dopamine system are not accompanied by obvious impairments on the rotating rod or in the open field (Lu et al., 2009, Sedelis et al., 2001, Fleming et al., 2013). The normalization of the voluntary movement behaviour at later time points in the open field are suggestive of a specific motor initiation problem that is distinct from hypoactivity behaviour described for some models of Parkinson’s disease, when defects in the nigrostriatal system cause disturbance of the central motor circuit as well (Sedelis et al., 2001, Fleming et al., 2013).

The motor control system in mammals is organized hierarchically, involving both cortical and subcortical circuits. In wildtype mice Syap1 is strongly expressed in the cerebellum (Schmitt et al., 2016) and in cerebellar nuclei (Fig. 6), particularly in Purkinje cell somata and in the molecular layer, where numerous glutamatergic synapses are formed between Purkinje cells and parallel and climbing fibers (Schmitt et al., 2016). The specific motor deficit in *Syap1* mutant mice may reflect changes in complex cerebellar motor control functions (Thach, 2014, De Zeeuw and Ten Brinke, 2015). Purkinje cells are the only efferent output from the cerebellar cortex and exert their function through projections to the deep cerebellar nuclei. Among the cerebellar nuclei, the fastigial nucleus (FN) is the phylogenetically oldest nucleus and is involved in axial, proximal and ocular motor control (Manto et al., 2012, Zhang et al., 2016). The FN is considered to be one of the ultimate output systems for cerebellar function and modulates motor behaviour via the vestibulospinal and reticulospinal tracts (Zhang et al., 2016). Defects in this projection pathway may explain the loss of function in the accelerated and reversed rotarod. Little is known about the role of the FN in the mouse, but studies in other mammals suggest that the FN also contributes to movement initiation via a bi-synaptic projection from the FN to the primary motor cortex (Kelly and Strick, 2003). This might contribute to the atypical motor behaviour in early phases of the open field test. Based on our results, we suggest that cerebellar functions for fast adaptation to more complex movements and regulated motor initiation might be affected by deletion of Syap1 in mice. In wildtype mice, Syap1 is also found in cortical areas involved in movement control (Schmitt et al., 2016). Therefore we cannot exclude the possibility that the distinct motor behaviour in *Syap1* mutants is due to reduced capabilities to initiate motor behaviour or to a reduced internal motivation to explore the open field arena. Both functions are controlled by neural circuits that include dorsal or ventral striatal information processing (Grillner et al., 2005, Bostan and Strick, 2018). However, considering the overall phenotype, this possibility seems less likely, because the explorative behaviour in late phases of the open field is normal. Furthermore, anxiety-like behaviour and cognitive information processing in the NOR task and Pavlovian fear conditioning tasks, are almost normal. Thus, Syap1-deficiency does not fundamentally disturb information processing in cortical areas, the hippocampus or the amygdala.

**Figure 6:**
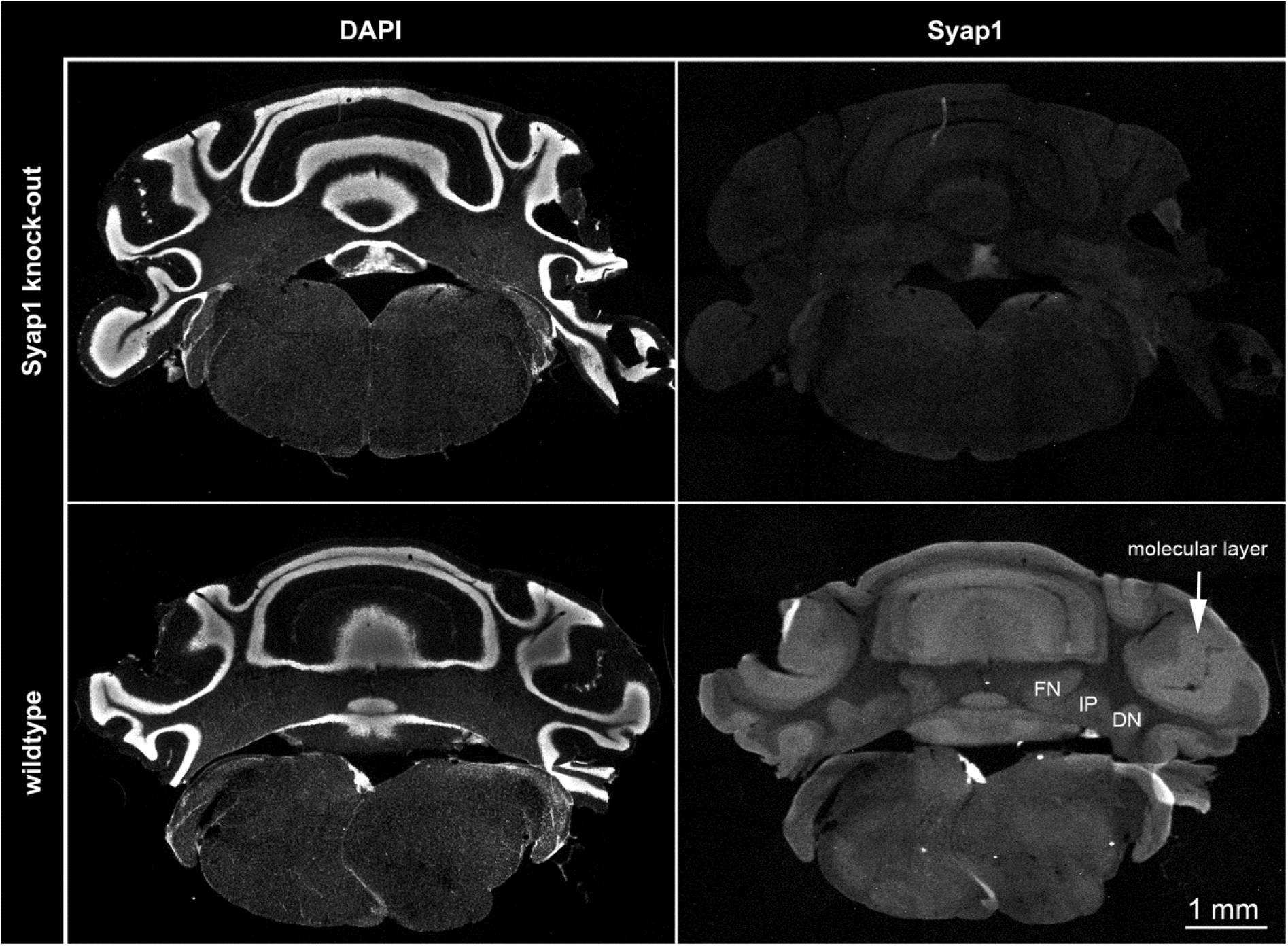
Anti-Syap1 immunoreactivity in the cerebellum. The images in the right panels show the immunoflourescence labelling of Syap1 protein in *Syap1* knockout (upper panels) and wildtype (lower panels) coronal sections of the cerebellum. DAPI labelling (left panels) was used as counterstain. Immunolabeling and image acquisition was performed as described earlier by Schmitt et al. (2016). The image illustrates the broad distribution of Syap1 protein in the cerebellum. High-resolution confocal analysis of the distribution of Syap1 in the cerebellar cortex has already been described earlier (Schmitt et al. 2016). The aim of these overview microscopy images is to illustrate the abundance of Syap1 in the deep cerebellar nuclei (fastigial nucleus (FN), interposed nucleus (IP), and dentate nucleus (DN). ML, molecular layer.

In *Drosophila*, deletion of the *Sap47* gene causes a ∼50 % decrease in larval learning scores for odor-tastant association (Saumweber et al., 2011), but motor performance of adult mutant flies has yet to be analysed. The lack of obvious defects of *Syap1* knockout mice in associative fear conditioning and novel object recognition, accompanied by a distinct motor skill deficit, does not necessarily point to a different molecular and cellular function of the evolutionarily conserved Sap47/Syap1 proteins, but might reflect a similar protein function expressed in the context of fundamentally different neural circuits.

In two human patients with autism spectrum disorder, copy number variants affecting the *SYAP1* gene have been observed (Prasad et al., 2012). At present, it is not clear whether SYAP1 dysfunction in humans causes any phenotype. The motor defects observed in the *Syap1* knockout mouse could be relevant for the discovery of such an association.

## Conclusion

We conclude that advanced motor skills, but not basic motor performance, depend on Syap1. Since it has been shown that Syap1 is highly concentrated in synaptic regions of cerebellar Purkinje cells (Schmitt et al., 2016), we suggest that there may be a causative link between cerebellar Syap1 expression and intact sensorimotor control. Basal ganglia and the cerebellum are interconnected at the subcortical level (Bostan and Strick, 2018). Therefore, we assume that the cerebellum and the dorsal striatum might represent target structures for a detailed analysis which could lead to a better understanding of the molecular and cellular function of Syap1.

## Materials and Methods

### Animals

All animal experiments were carried out in accordance with European regulations on animal experimentation and protocols were approved by the local authorities. Mice were housed individually and kept at a 12-hour dark-light cycle with access to food and water *ad libitum*. The cages (Tecniplast, 1264C Eurostandard Typ II, 267 × 207 × 140 mm) were kept in a Scantainer (Scanbur Ltd. Denmark) assuring stable conditions by maintaining a temperature of about 21°C and air humidity of about 55% through a constant air flow. Generation and verification of the *Syap1* null allele has been described before (Schmitt et al., 2016). Twelve knockout males (Y/-) and 12 wildtype littermates (Y/+) aged between 11 and 19 weeks were used for the experiments which were carried out during the light phase.

### Modified SHIRPA

Mice were individually placed in a viewing jar, a hollow acrylic glass cylinder of 18.7 cm height and 14.2 cm diameter, to observe and note down the following features: body position, tremor, palpebral closure, coat appearance, whisker appearance, lacrimation, defecation and urination. The next day, mice were placed in an arena (Tecniplast, 1284L Eurostandard Typ II L, 365 × 207 × 140 mm) and the following aspects were observed in the arena: transfer arousal, locomotor activity, gait, tail elevation, touch escape. The following features were also evaluated: skin colour, trunk curl, limp grasping, pinna reflex, corneal reflex, evidence of biting and vocalization.

### Open Field

A white square box made of frosted plastic (48 × 48 cm, height 50 cm), evenly illuminated with ca. 40 lux was used as open field arena. Mice were placed in the middle of the arena and were monitored for 30 min using a web cam-based system (camera: Logitech). Animal movements were tracked and analysed with the Video Mot Software (TSE, Germany). For analysis, the floor of the box was divided into different fields of interest. The following parameters were measured and compared between the centre of the arena (24 × 24 cm) versus the periphery: total distance travelled over time, travelling speed, time spent in the centre or periphery of the arena, number of entries to the centre and into the periphery.

### Rotarod

Motor skills were also analysed on the rotarod (Ugo Basile). In a first test phase, mice were investigated at a continuous speed of 5 rpm for 5 min. In a second test phase, the rotarod was set to accelerate from 5 rpm to ∼ 50 rpm within 5 min. In a third test phase (reverse rocking), mice were placed on the rod programmed to alternate rotating forwards and backwards to a final speed of 5 rpm. Accelerated and rocking rotarod were each performed on two subsequent days. For all test phases, the latency to fall off the rod was measured in seconds.

### Grip Strength

Grip strength of front and hind paws was measured as the obtained tension peak in Newton with a digital force gauge (Chattilon Digital Force DFI2). The procedure was repeated four times to acquire a mean value of grip strength.

### Novel object recognition (NOR)

The novel object recognition test was carried out in the open field arena (Leger et al., 2013). Two objects were presented: a cell culture flask (T75 Greiner, height about 16 cm) filled with water to the top and a 14.6 cm tower built of Lego bricks. Objects were placed in two diagonally opposing corners of the box 12 cm from each wall. During the first day, two identical objects were presented in a randomized fashion. Mice were observed and tracked for 10 min using the Video Mot Software (TSE Germany, camera: Logitech). The time the mice spent with their head in the fields of interest (2 cm distance surrounding each object) was measured manually. On the second day, mice were confronted with a familiar and a novel object. Here, a triplicate of the objects was used to avoid olfactory cues or influence. Position of objects was randomized to avoid position bias.

### Pavlovian Fear Conditioning

For Pavlovian fear conditioning, the Multi Conditioning System from TSE (256060 series) was used. The animals were monitored and tracked with the TSE MCS FCS – SQ MED software. On the first day, mice were placed in Context A, a square clear acrylic box placed on a metal grid. After a habituation time of 60 s a tone was presented three times (CS, intensity 85 dB SPL, 10000 Hz) that lasted 10 s with an inter stimulus interval (ISI) of 20 s. The CS was accompanied by a foot shock of 0.7 mA (US) which was delivered via the electric grid during the last second of tone presentation. After an additional time of 30 seconds, mice were transferred back to their home cage. To test for cue memory the next day, mice were placed in Context B, a clear acrylic cylinder placed on black, rough plastic. After an initial time of 60 s, the CS (tone) was delivered again for three times for 10 s with an ISI of 20 s, without administration of the US. On the third day, mice were placed back in the fear conditioning context (training Context A) for 6 min without CS presentation to recall contextual memory.

### Dark-Light Transition

The dark-light transition test was performed in the open field arena. For this purpose, a red acrylic glass box of 47 × 16 × 25 cm was positioned in the box covering approximately one third of the arena with a square opening serving as an entrance for the mouse to get into the dark compartment. Mice were placed in the light compartment and their movements were tracked for 10 min using the Video Mot Software (TSE Germany, camera: Logitech). The following parameters were recorded and analysed: distance travelled in the light compartment and time spent in each compartment.

### Elevated Plus Maze

The elevated plus maze consisted of a cross with two closed and two open arms made of white frosted plastic (TSE Germany, length of arm: 30 cm, width: 5 cm, height of closed arm: 15 cm, height above ground: 48 cm, luminosity adjusted to ca. 60 lux). Mice were placed on one of the open arms and their movement was tracked for 10 min using the Video Mot Software (TSE Germany, camera: Logitech). The following parameters were analysed and compared between open and closed arms: distance travelled and time spent in open and closed arms.

### Statistical Analysis

All statistical analyses were performed with GraphPad Prism 6. Data are presented as means with standard error of the means (± SEM). To account for statistical differences between genotypes and conditions depending on the experiment, two-way ANOVAs with post hoc Sidak’s multiple comparison test or unpaired *t*-tests were performed. Results were considered statistically significant at p < 0.05. A summary of the statistical results is given in supplementary table S1.

## Acknowledgements

We are grateful to Dr. Lill Andersen for essential contributions in generating the Syap1 knockout mouse and to Regine Sendtner and Michaela Keßler for excellent technical assistance.

## Author contributions

Conceptualization: M.S., R.B., E.B.; Methodology: C.RvC., D.S., R.B., T.R.; Formal analysis: C.RvC.; Investigation: C.RvC., D.S.; Resources: T.R., M.S.; Data curation: C.RvC., R.B.; Writing - original draft: C.RvC., R.B., E.B.; Writing - review &editing: M.S., R.B., E.B., T.R., C.RvC., D.S.; Supervision: M.S., R.B., E.B.; Funding acquisition: M.S., R.B., E.B.

